# Active Liposomal Targeting for Head and Neck Squamous Cell Carcinoma Treatment

**DOI:** 10.64898/2025.12.03.691158

**Authors:** Almudena del Campo-Balguerías, Carmen Martín-Hernández, Cristina Herrero-Igartua, Fernando de Andrés, Abdessamad Gueddari, María García-Bravo, Alberto Ocaña, Iván Bravo, Carlos Alonso-Moreno, Corina Lorz

**Affiliations:** Universidad de Castilla-La Mancha, Departamento de Química Inorgánica, Orgánica y Bioquímica-Centro de Innovación en Química Avanzada (ORFEO-CINQA), Unidad NanoDrug, Facultad de Farmacia, Ave Dr. Jose Maria Sánchez Ibáñez s/n, 02071 Albacete, Spain; Biomedical Innovation Unit, CIEMAT (ed 70A), Ave Complutense 40, 28040 Madrid, Spain; Research Institute 12 de Octubre i+12, University Hospital 12 de Octubre, Ave Córdoba s/n, 28041 Madrid, Spain; Universidad de Castilla-La Mancha, Departamento de Química Analítica y Tecnología de Alimentos, Facultad de Farmacia, Ave Dr. Jose Maria Sánchez Ibáñez s/n, 02071 Albacete, Spain. Instituto Regional de Investigación Científica Aplicada IRICA, Ave Camilo José Cela 1, 13005 Ciudad Real, Spain; Centro de Investigación Biomédica en Red de Enfermedades Raras (CIBERER), Ave Monforte de Lemos 3-5, 28029 Madrid, Spain; Instituto de Investigación Sanitaria Fundación Jiménez Díaz (IIS-FJD, UAM), Ave Reyes Católicos 2, 28040 Madrid, Spain; Experimental Therapeutics in Cancer Unit, Instituto de Investigación Sanitaria San Carlos (IdISSC), Profesor Martín Lagos s/n, 28040 Madrid, Spain; START Madrid-FJD, Hospital Fundación Jiménez Díaz (FJD) Early Phase Program, Fundación Jiménez Díaz Hospital, Ave Reyes Católicos 2, 28040 Madrid, Spain; Centro de Investigación Biomédica en Red de Cáncer (CIBERONC), Ave Monforte de Lemos 3-5, 28029 Madrid, Spain; Universidad de Castilla-La Mancha, Departamento de Química Física, Unidad NanoDrug, Facultad de Farmacia, Ave Dr. Jose Maria Sánchez Ibáñez s/n, 02071 Albacete, Spain

**Keywords:** . head and neck cancer, liposomes, therapy, cisplatin, alpelisib, active targeting

## Abstract

Head and neck squamous cell carcinoma (HNSCC) is a major oncological challenge due to its high recurrence rates, biological heterogeneity, and poor long-term survival outcomes. Despite advances in surgery and radiotherapy, chemotherapeutic options are limited. Cisplatin (CDDP) remains the mainstay, while small molecule inhibitors, such as the selective PI3Kα inhibitor Alpelisib, are still under clinical evaluation. However, systemic side effects and suboptimal tumor specificity hamper the clinical utility of both standard and novel specific therapies.

In this study, we developed passive- and active-targeting liposomes encapsulating Cisplatin (CDDP) and Alpelisib. To achieve active targeting, the anti-epidermal growth factor receptor (EGFR) antibody Cetuximab was conjugated to the liposome surface. All formulations were characterized based on size, morphology, encapsulation efficiency, and drug release profiles. *In vitro* assays demonstrated that liposomal encapsulation preserved the cytotoxic effects of both CDDP and Alpelisib in HNSCC cell lines, with CDDP showing adequate encapsulation efficiency for *in vivo* evaluation.

*In vivo* near-infrared fluorescence imaging demonstrated that active-targeting liposomes achieved greater tumor accumulation and a better tumor-to-spleen ratio compared to passive formulations. Tumor growth by CDDP-loaded active-targeting liposomes was comparable to that of the free drug, likely due to the hydrophilic nature and rapid release of CDDP from liposomes in physiological conditions. Importantly, blood chemistry analysis indicated a favorable toxicity profile for the targeted liposomal formulation.

These findings highlight the potential of liposome-based targeted strategies to improve therapeutic outcomes in HNSCC and underscore the value of nanotechnology in the development of next-generation oncological therapies.

## 1. Introduction

Head and neck cancer (HNC) accounts for approximately 4.5% of global cancer incidence and mortality [1,2]. HNC includes malignancies that originate in the upper aerodigestive tract, including the lips, oral cavity, salivary glands, larynx, nasopharynx, hypopharynx, and oropharynx. The most common subtype of HNC is head and neck squamous cell carcinoma (HNSCC), which originates from the squamous cells found on epithelial surfaces of these anatomical regions [3,4]. A high recurrence rate is typical of HNSCC, occurring in about one-third of cases. Patients with recurrent or metastatic disease often have a poor prognosis, with survival typically less than one year after diagnosis [5]. Moreover, HNSCC is a highly heterogeneous disease, with differences in molecular and clinical behavior depending on tumor location and causative factors, such as exposure to carcinogens and human papillomavirus infection [3]. Given the aggressive nature and biological complexity of this tumor, there is a pressing need for more effective treatment strategies.

Current treatment of HNSCC involves surgery and radiation therapy (RT), either alone or in various combinations that include chemotherapy. The specific approach depends on the stage and the primary location of the tumor [6]. For decades, Cisplatin (CDDP) combined with RT has been the standard of care for patients with locally advanced disease. CDDP-based combination treatments improve overall survival and progression-free survival; however, patients treated with CDDP experience more side effects compared to those who received radiation alone. These side effects include hematologic toxicities, mucositis, and nausea/vomiting [7]. Consequently, despite the favorable outcomes and extensive use of CDDP, its toxicity limits its clinical utility in the treatment of HNSCC, underscoring the necessity for less toxic approaches [8].

Despite the advances in surgery and RT techniques, the lack of significant improvement in the overall survival of patients with advanced HNSCC has driven the development of novel approaches, including molecularly targeted therapies, such as Cetuximab, an anti-epidermal growth factor receptor (EGFR) antibody, and immunotherapies, including the immune checkpoint inhibitors Pembrolizumab and Nivolumab. [8]. Among the signaling pathways identified as actionable therapeutic targets in HNSCC, the PI3K/AKT/mTOR pathway controls numerous cellular processes, including cell growth, survival, and metabolism. Its dysregulation is a well-established hallmark of cancer. More than 90% of HNSCC tumors show activation of this oncogenic signaling pathway through various mechanisms, including EGFR activation, PI3K overexpression, and amplification of the PIK3CA gene (which encodes the catalytic subunit of the PI3K kinase), among others [9,10]. Targeting the PI3K/AKT/mTOR pathway with specific inhibitors represents a promising therapeutic strategy for HNSCC. Alpelisib (BYL719), a selective PI3Kα inhibitor approved by the FDA for the treatment of PIK3CA-mutant metastatic breast cancer [11], is one such inhibitor. This drug is in clinical assessment for the treatment of HNSCC alone or in combination with other drugs (i.e.: NCT03138070; NCT01602315; NCT04997902). Although Alpelisib has a more favorable toxicity profile compared to pan-class PI3K inhibitors [12], its clinical use remains associated with dose-limiting toxicities and a high incidence of treatment-related adverse events. Common side effects include hyperglycemia, nausea, loss of appetite, diarrhea, vomiting, and induction of diabetic ketoacidosis [13–15].

Drug delivery systems (DDS) offer a valuable strategy for cancer treatment, enabling the use of effective but highly toxic drugs by improving their delivery and reducing systemic side effects [16,17]. Among the various materials employed for DDS formulation, liposomes are the most extensively studied and stand out for their promising properties as therapeutic carriers [18]. Their unique advantages include high biocompatibility, biodegradability, and low toxicity. Liposomes enable targeted delivery, enhance drug stability, and provide controlled release, thereby improving the therapeutic index and bioavailability of encapsulated agents. Liposomes have been approved by the FDA for various treatments, underscoring their importance in modern nanomedicine [19,20].

Nanotechnology can facilitate precise targeting in cancer therapy beyond the encapsulation strategy employed to mitigate adverse effects and regulate the release of therapeutic agents. Two mechanisms enhance nanoparticle (NP) accumulation in tumors: passive and active tumor targeting. Tailored design of NP-based DSS can achieve one or both. Passive targeting exploits the enhanced permeability and retention (EPR) effect, enabling NPs to accumulate at tumor sites based on their leaky vasculature [21,22]. Active targeting involves functionalizing NP surfaces with ligands, antibodies, peptides, or biomacromolecules that recognize tumor-specific receptors, enhancing their specificity toward tumor cells [23]. One such receptor, EGFR, is a key target in cancer therapy due to its involvement in cell proliferation, apoptosis, angiogenesis, and metastasis. Its overexpression, observed in approximately 90% of HNSCC cases, is associated with poor prognosis [24,25]. Active targeting strategies that modify the NP surface with the anti-EGFR antibody Cetuximab have shown promise in enhancing nanoparticle selectivity [26–29].

Previous studies have reported successful encapsulation of CDDP into liposomes for passive targeting in HNSCC [30–32]. For example, Harrington *et al.* demonstrated that PEGylated liposome-encapsulated CDDP enhanced tumor radiosensitivity and reduced nephrotoxicity and systemic side effects when combined with radiotherapy compared to free CDDP [32]. Lan *et al.* encapsulated CDDP alone and in combination with an anti-programmed cell death protein-1 (PD-1) antibody into liposomes and delivered them in dissolvable microneedles. These microneedles were released locally from the skin after insertion. Both approaches achieved greater tumor weight and volume reduction than free CDDP, with minimal systemic toxicity [30,31]. However, there have been only a few reports of active-targeting liposomes encapsulating CDDP for the treatment of HNSCC. In 2015, Lukianova-Hleb *et al.* proposed a complex quadruplet-treatment strategy for HNSCC involving an actively targeted liposomal formulation of CDDP, gold nanoparticles conjugated with antibodies, and radiotherapy [33]. A study by Yang *et al.* developed a more straightforward nanotherapeutic approach based on folic acid-functionalized liposomes co-encapsulating CDDP and paclitaxel for active targeting in HNSCC. This strategy resulted in a significant reduction in tumor size compared to other tested formulations, highlighting the therapeutic advantages of targeted co-delivery systems [34].

In this proof-of-concept study, we developed both passive- and active-targeting liposomal formulations of CDDP (conventional chemotherapy) and Alpelisib (targeted therapy) to improve their therapeutic efficacy in HNSCC treatment. Active targeting was achieved by functionalization of drug-loaded liposomes with Cetuximab, enabling selective binding to EGFR-overexpressing tumor cells. We conducted *in vitro* and *in vivo* preclinical studies to evaluate their therapeutic potential. These DDS aim to enhance intracellular drug tumor uptake and tumor penetration while reducing systemic toxicity and off-target effects on healthy tissues. This study significantly advances the development of targeted nanotherapy for HNSCC by demonstrating the feasibility and effectiveness of vectorized, liposome-based drug delivery systems, thereby paving the way for future research in nanotechnology-driven HNSCC treatment. Notably, this is the first report on the encapsulation of Alpelisib in liposomes for HNSCC therapy.

## 2. Materials and methods

### 2.1. Chemicals

1-ethyl-3-(-3-dimethylaminopropyl) carbodiimide hydrochloride (EDC) (99.95% purity), Cisplatin (CDDP) (99.84% purity) and Alpelisib (BYL719) (purity ≥ 99.9%) were purchased from MedChemExpress (Monmouth Junction, NJ, USA). N-Hydroxysuccinimide (NHS) (purity ≥ 98%), Pluronic® F-127, DSPE-PEG (1,2-distearoyl-sn-glycero-3-phosphoethanolamine-N- [amino(polyethylene glycol)]), L-α-phosphatidylcholine from soybean (Lecithin) (purity > 95%) and dimethyl sulfoxide (DMSO) (purity ≥ 99.9%) were obtained from Sigma-Merck (Darmstadt, Germany). DiR (purity > 99%) was purchased from Biotium (Freemont, CA, USA). Cetuximab was from Merck (Darmstadt, Germany). Dichloromethane (DCM) and acetone solvents were purchased from Análisis Vínicos (Tomelloso, Spain). Decanoyl and octanoyl glycerides (glycerides) were acquired from ChemoSapiens (Barcelona, Spain). Dialysis membranes tubes (Spectra/Por™, MWCO 12-14 Da, flat width: 25 mm) were obtained from Dilabo (Barcelona, Spain). Dulbecco’s Phosphate Buffered Saline (PBS 1×) was purchased from LabClinic (Barcelona, Spain).

### 2.2. Formulation of liposomes

#### 2.2.1. Formulation of non-conjugated liposomes loaded with Alpelisib, CDDP, DiR, or DiD

Loaded liposomes were prepared using an optimized solvent-injection method [35–37]. Briefly, 10 mL of Pluronic® F-127 (1% w/v in Milli-Q® water) was added to a round-bottom flask to serve as the aqueous phase. The organic phase was prepared by mixing 500 µL of soy lecithin (50 mg/mL in DCM), 70 µL of glycerides, and 1 mg of either Alpelisib, DiD, or DiR (5 µL of a 200 mg/mL solution in DMSO), using a vortex mixer. The organic mixture was then added to the aqueous phase and mixed under high-speed homogenization at 14,000 rpm for 10 minutes, forming O/W emulsion. Finally, the organic solvent was evaporated at 40 °C, yielding the corresponding liposomal formulations. For the CDDP-loaded liposomes, 2 mg of the drug powder was incorporated into the aqueous phase before the addition of the organic phase.

Subsequently, the liposomes were washed by dialysis in phosphate-buffered saline (PBS, 1M, PH 7.4) using dialysis membrane tubes with a molecular weight cut-off (MWCO) of 12–14 kDa. The liposomal formulations were then transferred into the dialysis membranes and immersed in PBS at a 1:10 volume ratio, followed by magnetic stirring for 20 minutes. All Alpelisib-, DiR-, and DiD-loaded liposomes were obtained at a theoretical concentration of 0.1mg/ml. The theoretical concentration of CDDP-loaded liposomes was 0.2 mg/ml.

#### 2.2.2. Formulation of Cetuximab-conjugated liposomes loaded with Alpelisib, CDDP, DiR, or DiD

First, the loaded liposomes were prepared following the same protocol described in the previous section, with the addition of 50 µL of a DSPE-PEG solution (concentration of 20 mg/mL in MeOH) to the organic phase. For antibody conjugation, 250 µL of Cetuximab (20 µg/mL in 0.1 M PBS, pH 7.4), 160 mg of EDC, and 40 mg of NHS were added to 0.5 mL of PBS (0.1 M, pH 5.8) and stirred for 1 hour at room temperature. Subsequently, 400 µL of DSPE-PEG-coated liposomes were added to the mixture and left under stirring for 3 hours at room temperature. Finally, Cetuximab-conjugated liposomes were washed by dialysis in PBS (0.1M, pH 7.4).

#### 2.2.3. Formulation of conjugated and non-conjugated CDDP-loaded liposomes for in vivo tumor growth inhibition assay

For the *in vivo* experiments, the concentration of encapsulated CDDP was increased to achieve the required therapeutic doses. To this end, CDDP-loaded liposomes were formulated following the same protocol previously described, with the modification that 7.5 mg of CDDP powder was weighed and added to 5 mL of Pluronic® F-127 (1% w/v in Milli-Q® water) as the aqueous phase. The organic phase was not altered, and the same reagent volumes were used as in the original method. Cetuximab conjugation was carried out following the same procedure described above. After that, liposomes were washed by dialysis in PBS (0.1 M, pH 7.4) following the same procedure.

### 2.3. Physicochemical characterization and stability of liposomes

The size, polydispersity index (PdI), and Z-potential of the liposomes were analyzed using dynamic light scattering (DLS) on a Zetasizer Nano ZS instrument (Malvern Instruments, Malvern, UK). Data were processed using the multimodal number distribution software provided with the instrument. DLS measurements were used to monitor the hydrodynamic radius (R_H_) and PdI of the liposomal formulations over four weeks.

All liposome formulations were stored at 4 °C and subsequently incubated in PBS at 37 °C before DLS analysis, which was used to assess their average size (nm) and PdI.

The morphology of the nanoparticles was characterized by transmission electron microscopy (TEM). High-resolution images were acquired using a JEOL JEM-2100 microscope (Jeol Ltd., Japan) operated at 200 kV. The microscope was equipped with an Oxford Instruments EDS detector for elemental analysis. For sample preparation, nanoparticle suspensions were diluted in distilled water, deposited onto copper grids and dried under ambient conditions. To minimize electron beam-induced damage, images were acquired under low-dose conditions. Image analysis was carried out using Digital Micrograph™ software from Gatan

### 2.4. Encapsulation efficiency (EE%)

For liposomal NPs, the EE% of the conjugated and non-conjugated formulations was determined by dialysis in PBS. For this procedure, the liposomal formulations were transferred into the dialysis membranes and immersed in Milli-Q® water at a 1:5 volume ratio. The amount of free (non-encapsulated) drug in the supernatant, required to calculate the mass of encapsulated drug (in mg), was quantified using different techniques depending on the compound.

Alpelisib content was quantified using an HPLC system. Briefly, Alpelisib containing supernatant solutions were 1:2 diluted in the selected mobile phase, and the analyte was baseline separated in less than 1.4 min under isocratic elution using H2O:acetonitrile (20:80) as mobile phase at 45°C and then determined by mass spectrometry in positive ion mode under the following conditions: drying gas flow of 12 L/min, nebulizer pressure of 2.41×10 Pa, drying gas temperature of 300°C, and capillary voltage of 5000 V. Quantification was achieved using the external calibration method and single ion monitoring (SIM) for the selected analyte. Consequently, the 442.7 mass-to-charge (m/z) ratio was monitored. Initial identification was performed using full scan mode, and both the mass spectrum and retention time (0.8 min) were used to confirm the presence of Alpelisib. All experiments were performed in triplicate, and the results are expressed as the mean ± standard deviation.

CDDP was quantified using inductively coupled plasma mass spectrometry (ICP-MS). CDDP concentrations were thus determined using flow injection analysis under these conditions: RF power 1550 W; RF matching 1.80 V; carrier-gas flow 0.99 L/min, no make-up gas; sampling depth 10 mm; nebulizer pump rate 0.1 rps; spray chamber temperature 2°C. The ICP−MS instrument was operated in helium collision mode for unsurpassed interference removal. A typical performance test in He mode is as follows: He flux 5 mL/min, m/z 7 (465 cps), m/z 89 (2616 cps), m/z 205 (1012 cps). Every sample contained 1 μg/L erbium as internal standard for ICP−MS measurements and all determinations were conducted by monitoring the 195Pt signal. The ICP−MS instrument was tuned using a solution containing 1 μg/L each of Ce, Co, Li, Mg, Tl and Y (Agilent), and calibration curves were obtained using aqueous standard solutions in identical matrix to the samples (with regard to internal standard and 2% nitric acid) with appropriate stock standards dilutions. All experiments were performed in triplicate.

Finally, DiR and DiD content was determined by UV-Vis spectrophotometry. Absorbance was measured at 275 nm using a Cary 100 spectrophotometer at room temperature, with a slit width of 0.4 nm and a scan rate of 600 nm/min. The following equation was used to calculate the EE%:

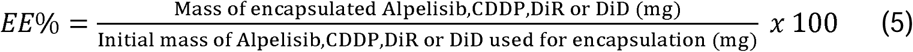

All experiments were conducted independently in triplicate, and results are presented as mean ± standard deviation.

### 2.5. Drug release studies

A volume of 2.5 ml of conjugated and non-conjugated drug-loaded liposomes was sealed in a dialysis membrane, which was suspended in 10 mL of phosphate-buffered saline (PBS, 0.1 M, pH 7.4) at 37°C under constant magnetic stirring. At regular time intervals, 1 mL of the external medium was withdrawn and replaced with an equal volume of fresh PBS to maintain sink conditions. The concentration of the released drug in each sample was quantified using the analytical techniques previously described in the encapsulation and loading efficiency section. All experiments were performed in triplicate, and the results are expressed as the mean ± standard deviation.

### 2.6. Antibody-conjugation quantification

Cetuximab-conjugated liposomes were separated from unbound Cetuximab using a Sepharose CL-4B column and eluted with phosphate-buffered saline (PBS, pH 7.4) [38–40]. Following separation, the concentration of non-conjugated Cetuximab in the elution was quantified using the bicinchoninic acid (BCA) assay, following the standard protocol [41,42]. For this, the sample was incubated with BCA reagent in a 96-well plate for 30 minutes in the dark. Subsequently, the sample’s absorbances were measured at 563 nm using a Multiskan™ GO Microplate Spectrophotometer (ThermoFisher Scientific, Finland) to determine the concentration of free (non-conjugated) Cetuximab.

### 2.7. Cell lines and cell culture

Human HNSCC-derived cell lines (Detroit562, FaDu) were obtained from the ATCC (Rockville, MD, USA) or kindly provided (Cal33, Cal27, WSU-HN6) by Dr. JS Gutkind (Department of Pharmacology and Moores Cancer Center, University of California, San Diego, La Jolla, CA, USA). Cells were cultured in Dulbecco’s Modified Eagle’s Medium (DMEM), supplemented with 10% (v/v) fetal bovine serum (FBS, Gibco), 1% (v/v) penicillin–streptomycin, and (Detroit562) with 1X non-essential amino acids (all from Gibco, Thermo Fisher Scientific, Walthan, MA, USA). Cells were maintained at 37 °C in an atmosphere of 5% CO2 and 95% humidity. Cell cultures were routinely tested for mycoplasma using a PCR-based detection kit (Venor™ GeM, Sigma-Aldrich).

### 2.8. Cell viability assays

Cell viability assays were performed using Cell Counting Kit-8 (CCK-8, Dojindo, Munich, Germany) which is based on the measurement of the metabolic cell activity. Cells (5,000-20,000 per well) were plated in 96-well plates for 24 h followed by incubation with escalating concentrations of free drug or liposome-loaded drug. After 72 h incubation with the inhibitors, absorbance values at 450 nm were recorded in a spectrophotometer plate reader Infinite 200Pro (Tecan, Switzerland). Background absorbance (culture medium without cells) was subtracted from all measurements, and the data were normalized as percentage of control. Each experiment was performed at least three times, and each concentration point was replicated three to six times within each experiment. The corresponding Inhibitory Concentration 50 (IC50) was calculated with GraphPad Prism5 (GraphPad Software, San Diego, CA, USA). This value is defined as the concentration of drug causing a decrease of 50% in cell viability as measured by the CCK-8 assay.

For cell survival analysis, free Alpelisib was suspended in DMSO and CDDP was suspended in dimethylformamide (DMF), to a stock concentration of 10 mg/mL (22.6 mM) and 3 mg/mL (10 mM), respectively.

### 2.9. Flow cytometry

To study liposome uptake, 500,000 cells in 0.5 mL of cell culture media, were incubated with different dilutions (1/50; 1/500; 1/1000) of LIP-DiD or ACLIP-DiD (DiD concentration in stock 0.068 mg/mL) for 30 min. Cells were immediately fixed in 4% saline formalin and analyzed using the 640Red 780_60-A laser of a BD LSR Fortessa cell analyzer with BD FACSDiva Software (BD Biosciences, Franklin Lakes, NJ, USA). Further data analysis was performed with FlowJo software (BD Biosciences). The fluorescent signal of cells incubated in the absence of fluorochrome was used to set the threshold (0% positive events).

To analyze EGFR protein cell surface expression, 500,000 cells were incubated with 0.5 µl of anti-EGFR(528)-APC antibody (clone AY13, Biolegend) and analyzed with the 640Red 670_30-A laser in a BD LSR Fortessa cell analyzer using BD FACSDiva Software. The fluorescent signal of cells incubated in the absence of fluorochrome was used as background fluorescence control.

### 2.10. Murine xenograft models for the analysis of liposome tumor homing and tumor growth

Murine xenograft mouse models were established by subcutaneous injection of HNSCC cell lines. Briefly, Cal33 and WSU-HN6 cells were trypsinized and suspended in a mixture (1:2) of DMEM 10% FBS:VitroGel (The Well Bioscience, Monmouth Junction, NJ, USA). Five million cells (Cal33) or two million cells (WSU-HN6) were suspended in a total volume of 150 μl of the DMEM-Vitrogel mixture and subcutaneously injected in the right flank of 10-week-old immunocompromised nude (Foxn1^nu/nu^) female mice bred at CIEMAT’s facilities (animal house registration number: ES280790000183). After 3 weeks, all mice developed tumors. At this point, the mice were injected intravenously with 100 µl of DiR-labelled nanoparticles (≈ 6.4 µg DiR/mouse), and *ex vivo* fluorescence in the tumors was analyzed 48 hours later using an *in vivo* imaging system (IVIS Lumina System, Perkin Elmer, Waltham, MS, USA). Liposomal preparations were sonicated for 10 min at 4 °C before injection.

For tumor growth studies, Cal33 cells were injected into both flanks. When the average tumor volume reached approximately 100-150 mm3 (day eight after cell injection), mice were randomized into three groups (n = 8 tumors per group). Mice were treated with 4 mg/kg CDDP twice a week. Free CDDP was prepared fresh for each use by diluting a 10 mg/mL CDDP DMF stock solution in saline solution (final concentration of DMF 5.4%). ACLIP-CDDP was prepared as described above. Free CDDP and ACLIP-CDDP were administered intraperitoneally and intravenously, respectively. Tumor size was measured twice a week using a Vernier caliper and calculated using the formula: tumor size = tumor length × (tumor width)^2^ × 0.5. Blood samples (100 µL) from each mouse were collected at the end of the experiment and analyzed in an SMT-120V Veterinary Biochemistry Analyzer (Seamaty, Chengdu, Sichuan, China).

All animal experiments were conducted at CIEMAT’s facilities in compliance with Institutional Animal Care and Use Committee guidelines and approved by the Animal Welfare Department (reference: PROEX 045.8/21).

### 2.11. Immunohistochemical Analysis

For immunohistochemical (IHC) and hematoxylin-eosin (H&E) staining, 4μm thick formalin-fixed, paraffin-embedded tissue sections were deparaffinized and rehydrated through a graded ethanol series. For IHC, antigen retrieval was carried out in 10 mM Tris-EDTA buffer (pH 9.0) by microwave heating. Tissue permeabilization was achieved using 0.1% Triton X-100 in PBS for 10 min. Endogenous peroxidase activity was blocked by incubation in 3% hydrogen peroxide in methanol for 10 min. To reduce nonspecific binding, sections were incubated in 0.1% Tween-20 in PBS for 10 min, followed by blocking in 10% horse serum (HS) in PBS for 1 h.

Slides were then incubated overnight at 4 °C with a primary antibody against Ki67 (1:1000 dilution, #ab15580, Abcam, Cambridge, UK) diluted in 5% HS in PBS. Then, sections were incubated with a biotinylated anti-rabbit secondary antibody (1:4000 dilution, #711-065-152, Jackson ImmunoResearch, Cambridge, UK) in 5% HS in PBS for 1 h at RT. Signal amplification was performed using the ABC kit (Vector Laboratories, Newark, CA). Detection was carried out using DAB substrate (Vector Laboratories).

Stained slides were photographed at 200X magnification using a Leica DM2000 LED microscope and quantified using Fiji (ImageJ) software.

### 2.12. Statistical analysis of the data

Data are shown as mean ± standard error of the mean (SEM). Normality test were run to evaluate the distribution of the data. Unpaired t-test, with Welch’s correction when variances differed (F test), was used for pairwise comparisons. One-way ANOVA with Tukey’s correction test was used for multiple comparisons. * 0.001 ≤ pVal < 0.05; ** 0.0001 ≤ pVal < 0.001.

## 3. Results

### 3.1. Formulation and physicochemical characterization of liposomes

Non-loaded liposomes (LIP), Alpelisib- and CDDP-loaded liposomes (LIP-Alp and LIP-CDDP) were formulated following an optimized solvent injection method (Scheme 1A) [35–37]. For liposome formulation, acetone was used as a water-miscible organic solvent, while lecithin and glycerides served as sources of phospholipids and as the main components of the oil phase. The copolymer Pluronic was included to ensure the stabilization and emulsification of the nanodevices.

**Scheme 1.**
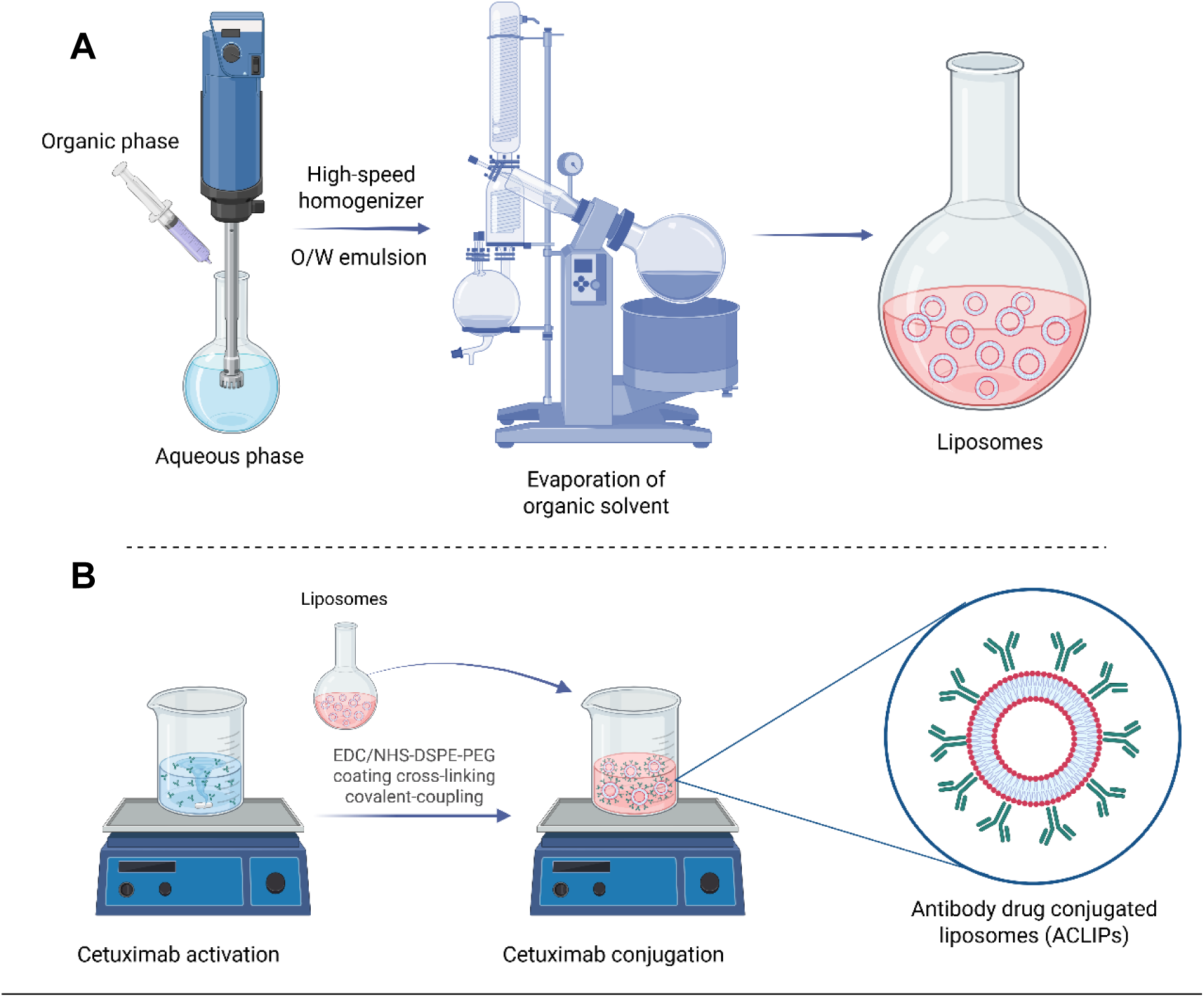
Schematic representation of the formulation of **(A)** LIP, **(B)** ACLIP.

For antibody conjugation, Cetuximab was covalently linked to a DSPE-PEG coating *via* chemical cross-linking, resulting in the formation of non-loaded Cetuximab-conjugated liposomes (ACLIP), Alpelisib-loaded Cetuximab-conjugated liposomes (ACLIP-Alp), CDDP-loaded Cetuximab-conjugated liposomes (ACLIP-CDDP), and DiR or DiD-loaded Cetuximab-conjugated liposomes (ACLIP-DiR, ACLIP-DiD) (Scheme 1B).

### 3.2. Characterization of the liposomal formulations

The characterization of LIPs and ACLIPs, both loaded and non-loaded, as well as Cetuximab-conjugated and non-conjugated, was carried out by dynamic light scattering (DLS) and transmission electron microscopy (TEM). Table 1 collects the physicochemical characteristics of the nanodevices. Encapsulation efficiency (EE%) was calculated according to established protocols [43]. The encapsulation of either Alpelisib or CDDP into the nanodevices did not significantly alter the physical properties of the formulations. As shown in Table 1, liposomes exhibited a comparable average hydrodynamic R_H_ of approximately 80 nm, with very low PdI (see Figure S1 in the Supplementary Material). Furthermore, no significant differences in size or polydispersity were observed between conjugated and non-conjugated liposomes. However, a shift in the surface charge of the liposomes, from approximately −20 mV to −10 mV, was observed following conjugation with Cetuximab, indicating a reduction in negative surface charges consistent with successful antibody coupling [27,44]. It is worth noting that the encapsulation efficiency (EE%) of all nanodevices, including the antibody-conjugated liposomes, was remarkably high, approaching 100%, which highlights one of the key advantages of liposomes as DDS.

**Table 1.** Characterization of LIP-CDDP, ACLIP-CDDP, LIP-Alp and ACLIP-Alp.

| Formulation | $R_H$ (nm) | Pdl | Z-Potential (mV) | EE% | Ab - conjugation (%) |
| --- | --- | --- | --- | --- | --- |
| **LIP** | 84,51 ± 3,94 | 0,09 ± 0,02 | -25,11 ± 1,51 | - | - |
| **ACLIP** | 74,32 ± 1,97 | 0,09 ± 0,01 | -13,26 ± 0,77 | - | 94 |
| **LIP-CDDP** | 81,32 ± 1,46 | 0,09 ± 0,09 | -20,72 ± 0,52 | 96,66 ± 0,70 | - |
| **ACLIP-CDDP** | 83,75 ± 3,27 | 0,13 ± 0,13 | -15,56 ± 1,24 | 98,28 ± 0,07 | 88 |
| **LIP-Alp** | 88,08 ± 1,36 | 0,09 ± 0,01 | -21,62 ± 0,65 | 92,07 ± 0,08 | - |
| **ACLIP-Alp** | 81,87 ± 1,36 | 0,06 ± 0,02 | -13,36 ± 1,88 | 89,30 ± 2,21 | 92 |

The antibody-functionalization process was based on carbodiimide chemistry, a widely used method due to its cost-effectiveness and high efficiency [45,46]. The carboxyl groups on Cetuximab were activated using the crosslinking agents EDC and NHS, enabling their conjugation to the primary amine groups present on the liposome surface following DSPE-PEG coating. The standard protocol of bicinchoninic acid assay (BCA) was followed to assess the efficacy of the conjugation, indicating a high degree of binding (see Figure S2 in the Supplementary Material) [41]. Representative TEM images of LIP (Figure 1A) and ACLIP (Figure 1B) show that both formulations exhibit a predominantly spherical morphology. The observed particle sizes and uniform distribution in TEM micrographs were consistent with the RH and PdI indices obtained by DLS measurements. As shown in Figure 1B, following conjugation with Cetuximab, the surface of the liposomes was altered, displaying an irregular core-shell morphology. This change may be indicative of successful antibody conjugation [41].

**Figure 1.**
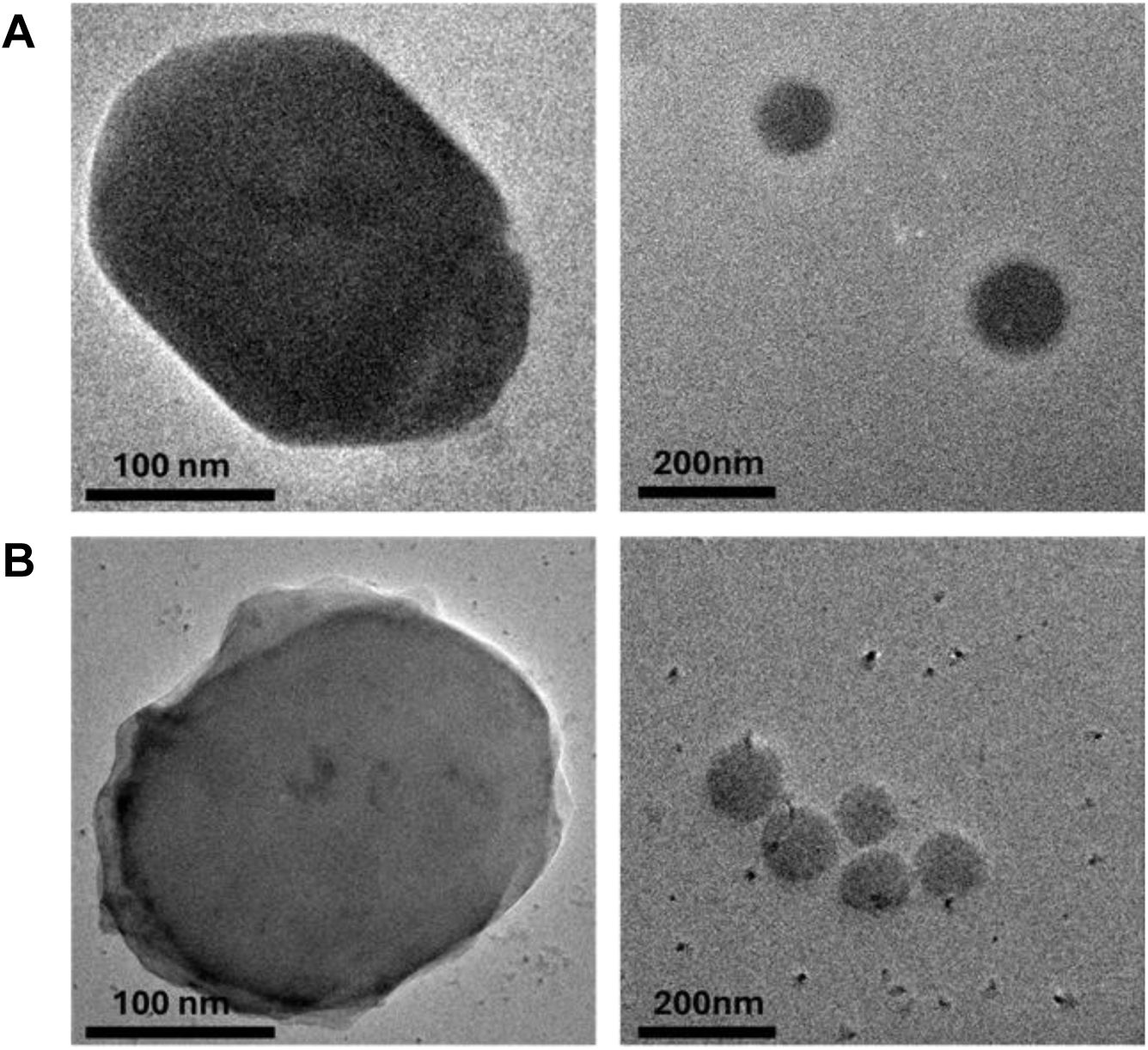
TEM images of **(A)** LIP and **(B)** ACLIP. Two different scale bars (100 nm and 200 nm) for each formulation.

### 3.3. Release profiles and storage stability of loaded liposomes

Figure 2A shows the colloidal stability of liposomal formulations, evaluated over a 30-day storage period in PBS at 4 °C by monitoring changes in R_H_ and PdI. The results showed minimal variations in particle size or polydispersity over the one-month observation period, indicating excellent formulation stability when stored at 4°C.

**Figure 2.**
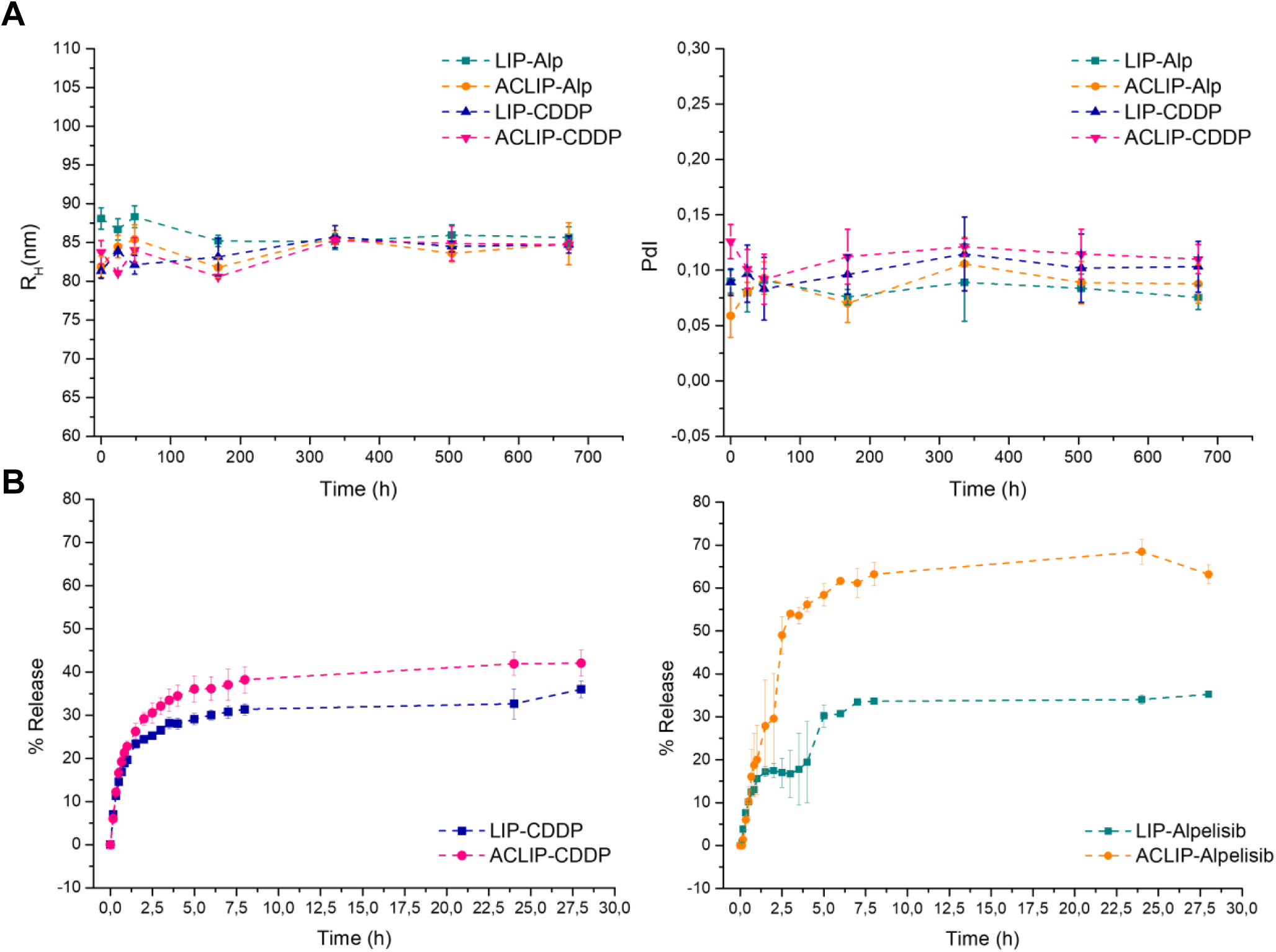
(A) Storage stability and release profiles of LIP-CDDP, ACLIP-CDDP, LIP-Alp, and ACLIP-Alp. Storage stability in PBS (pH 7.4) at 4°C. **(B)** I*n vitro* release profiles of the indicated formulation in PBS (pH 7.4) at 37°C. Data are presented as mean ± SEM (n= 3).

The drug release behavior of the nanodevices was then assessed using a dialysis method at pH 7.4 and 37 °C to simulate physiological conditions (Figure 2B) [47]. Formulations followed a triphasic release pattern with an initial rapid release phase within the first 2 hours. This initial burst is attributed to the desorption of drug molecules loosely associated with the liposome surface [48]. The burst release is more pronounced in CDDP compared to Alpelisip loaded liposomes, with cumulative CDDP release reaching 30%. This likely reflects the different physicochemical properties of the two drugs. CDDP, a small and hydrophilic molecule, readily undergoes aquation via chloride displacement in aqueous media, facilitating its release. In contrast, Alpelisib is a larger hydrophobic molecule, and different mechanisms may govern its diffusion through the liposomal bilayer. Following this phase, a more sustained release was observed for all formulations, remaining below 40% after 30 hours. Interestingly, ACLIP-Alp exhibited a significantly faster and more extensive release of Alpelisib compared to its non-conjugated counterpart, LIP-Alp. This enhanced release could be associated with the presence of Cetuximab on the nanoparticle surface, potentially affecting bilayer integrity and facilitating drug diffusion [48]. Despite differences in release kinetics, the antibody-conjugated and non-conjugated liposomes maintained controlled release profiles, reinforcing the potential of these systems for targeted drug delivery.

### 3.4. In vitro antiproliferative effect of free drug and drug-loaded liposomes

Alterations in the PIK3CA gene have been associated with sensitivity to Alpelisib in HNSCC and other cancer types [49]. Thus, we studied the effect of Alpelisib encapsulated in antibody-conjugated liposomes (ACLIP) or non-conjugated liposomes (LIP) in three HNSCC cell lines with different PIK3CA profiles. As previously reported [29,49], the PI3KCA mutant Cal33 cell line was the most sensitive to free Alpelisib, whereas FaDu (with PIKCA amplification) and Cal27 (wild type for PIK3CA) showed less sensitivity (Figure 3A). The antiproliferative effects of the encapsulated drug in both LIP and ACLIP were similar to the free drug, particularly in the Cal33 sensitive cell line, while ACLIP-Alp performed better than the free drug in the FaDu resistant cell line (Figure 3).

**Figure 3.**
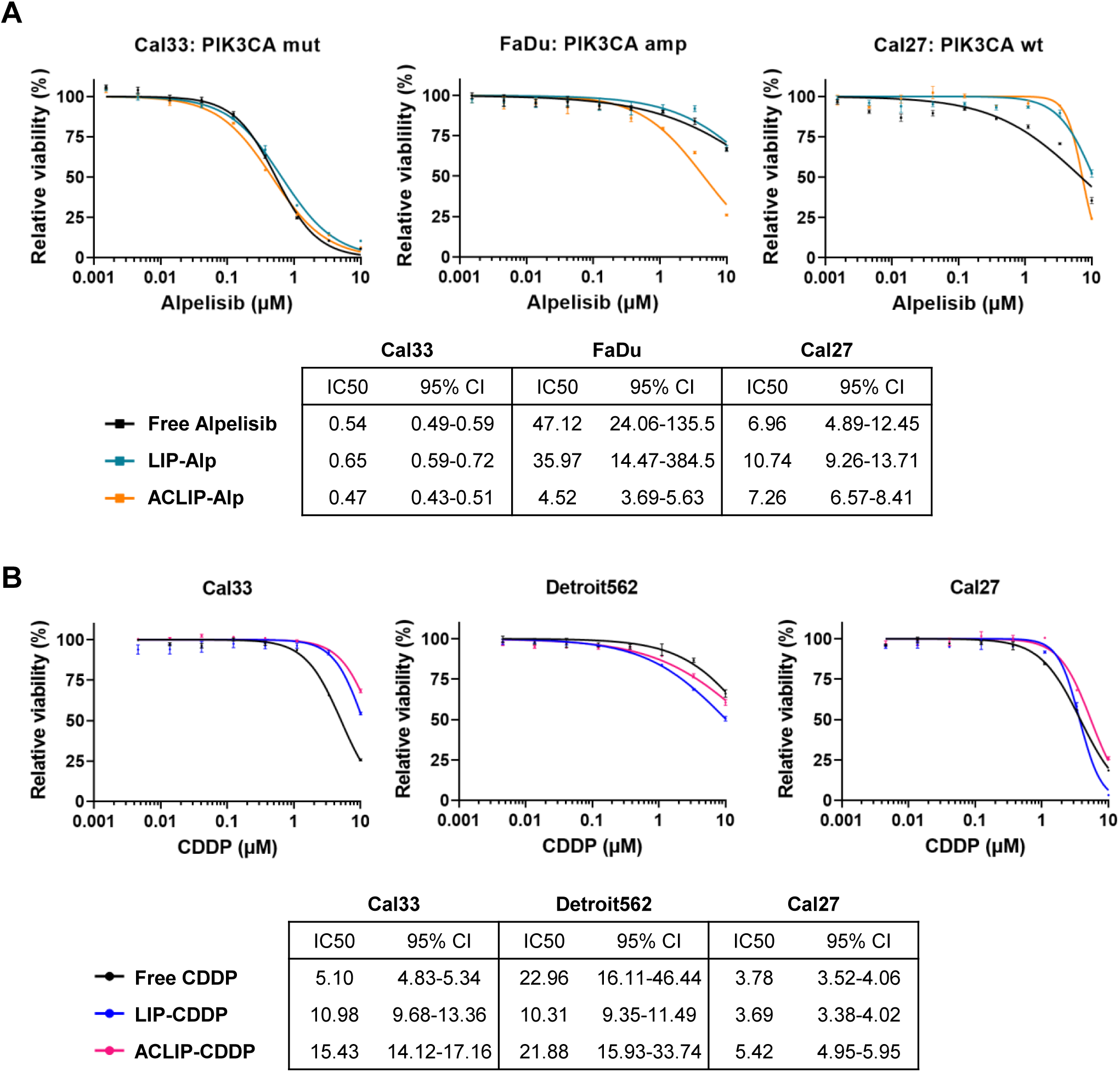
Cell viability of the indicated cell lines treated with increasing concentrations of free, LIP, and ACLIP-encapsulated Alpelisib **(A)** or CDDP **(B)**. The half-maximal inhibitory concentration 50 (IC50) for each cell line was determined using a non-linear regression fit of the data, represented by the curve in each graph. Data represent mean ± SEM (n = 3). The tables in **(A)** and **(B)** summarize the IC50 values (best-fit) and 95% CI values (profile likelihood) for the cell viability curves (log-inhibitor versus normalized response-variable slope) (n = 3). The data were analyzed using GraphPad Prism 9.5.1. CDDP: Cisplatin; LIP-Alp: liposome encapsulated Alpelisib; ACLIP-Alp: Cetuximab-conjugated liposome encapsulated Alpelisib; LIP-CDDP: liposome encapsulated Cisplatin; ACLIP-CDDP: Cetuximab-conjugated liposome encapsulated Cisplatin; IC50: inhibitory concentration 50; SEM: standard error of the mean; CI: confidence interval.

Next, we chose three HNSCC cell lines with varying sensitivity to free CDDP (Figure 3B). The encapsulated forms of CDDP showed comparable antiproliferative properties to the free counterpart (Figure 3B). Both empty LIP and ACLIP were non-toxic within the concentration range equivalent to that used to encapsulate the drugs (Figure S3 in the Supplementary Material). We have previously reported [29] that Cetuximab alone does not exhibit toxicity in HNSCC cell lines *in vitro* at the concentration used here.

Other studies have reported a similar potency for a free drug and its corresponding liposomal formulations [50]. This similarity may be attributed to the rapid cellular uptake of liposomes in an *in vitro* setting, followed by the release of the drug at a rate comparable to that of the free form. Flow cytometry analysis of HNSCC cells incubated with different concentrations of DiD-labeled liposomes confirmed their rapid internalization (≥ 98% of the cells were positive for DiD after 30 min incubation with the liposomes) regardless of whether the liposomes were conjugated with Cetuximab (Figure 4 and Figure S4 in Supplementary Material). Consistently, no significant differences were observed in the antiproliferative effects between drug-loaded LIP and ACLIP liposomes in our *in vitro* experiments (Figure 3). However, these experimental conditions do not fully replicate the complexity of the tumor microenvironment, where factors such as vascularization, immune response, and tissue-specific accumulation play crucial roles. To more effectively assess the benefits of passive targeting through the EPR effect and active targeting via Cetuximab-mediated delivery, we conducted *in vivo* experiments.

**Figure 4.**
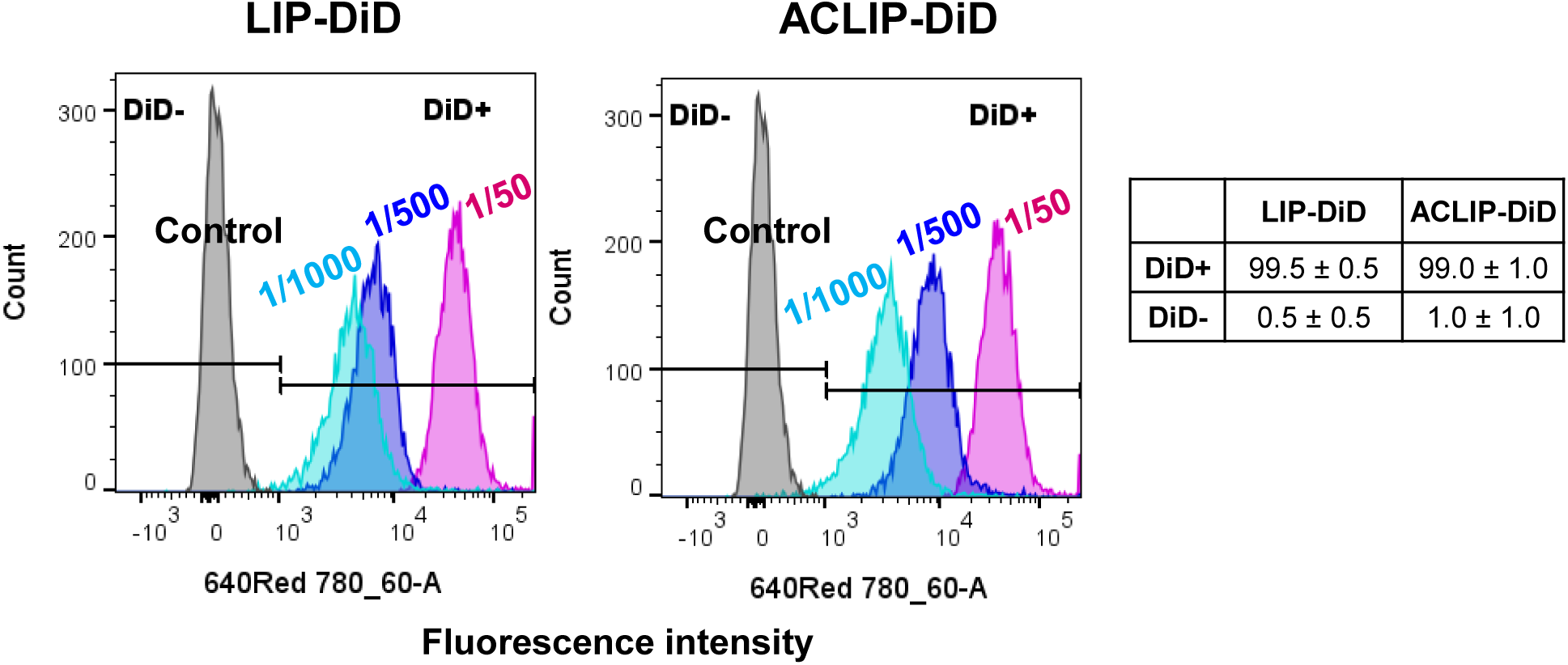
Overlay plots of flow cytometry histograms of Cal33 cells incubated in the presence of different dilutions of LIP-DiD or ACLIP-DiD for 30 min. Y axis: cell count; X axis: DiD fluorescence intensity. Background fluorescence in the absence of DiD-loaded liposomes is indicated by the grey peaks. Numbers in the table indicate mean ± SEM % of DiD positive and negative cells (n = 3). The graphs correspond to overlay plots of a representative experiment of at least three with similar results. LIP-DiD: liposome encapsulated DiD; ACLIP-DiD: Cetuximab-conjugated liposome encapsulated DiD.

### 3.5. In vivo tumor targeting of fluorescent-labeled liposomes in an EGFR-positive HNSCC xenograft model

To determine whether an active-targeting strategy (ACLIP) could lead to increased carrier accumulation at the tumor site compared to a non-targeted liposomal delivery system (LIP), we performed *ex vivo* imaging studies. We used a well-established xenograft tumor model [29,51,52] to evaluate the biodistribution and tumor-targeting potential of the near-infrared fluorescent DiR dye encapsulated into both types of liposomes. Far-red fluorochromes are particularly well-suited for this type of imaging studies due to their low background fluorescence in tissues at the appropriate excitation and emission wavelengths.

First, we analyzed the expression of our ACLIP target, EGFR, on the cell surface of three HNSCC cell lines using flow cytometry. We have previously shown that WSU-HN6, Cal33, and Detroit562 exhibit varying levels of EGFR copy number and total protein expression [51]. In our flow cytometry analysis, WSU-HN6 and Cal33 demonstrated higher median fluorescence for EGFR labeling on the cell surface compared to Detroit562 (Figure 5A). Consequently, we selected these two cell lines for the liposome tumor homing murine xenograft model (Figure 5B). Moreover, the tumors generated by these cell lines represent the extremes of HNSCC aggressiveness, with WSU-HN6 being undifferentiated and Cal33 being well-differentiated (Figure S5 in the Supplementary Material).

**Figure 5.**
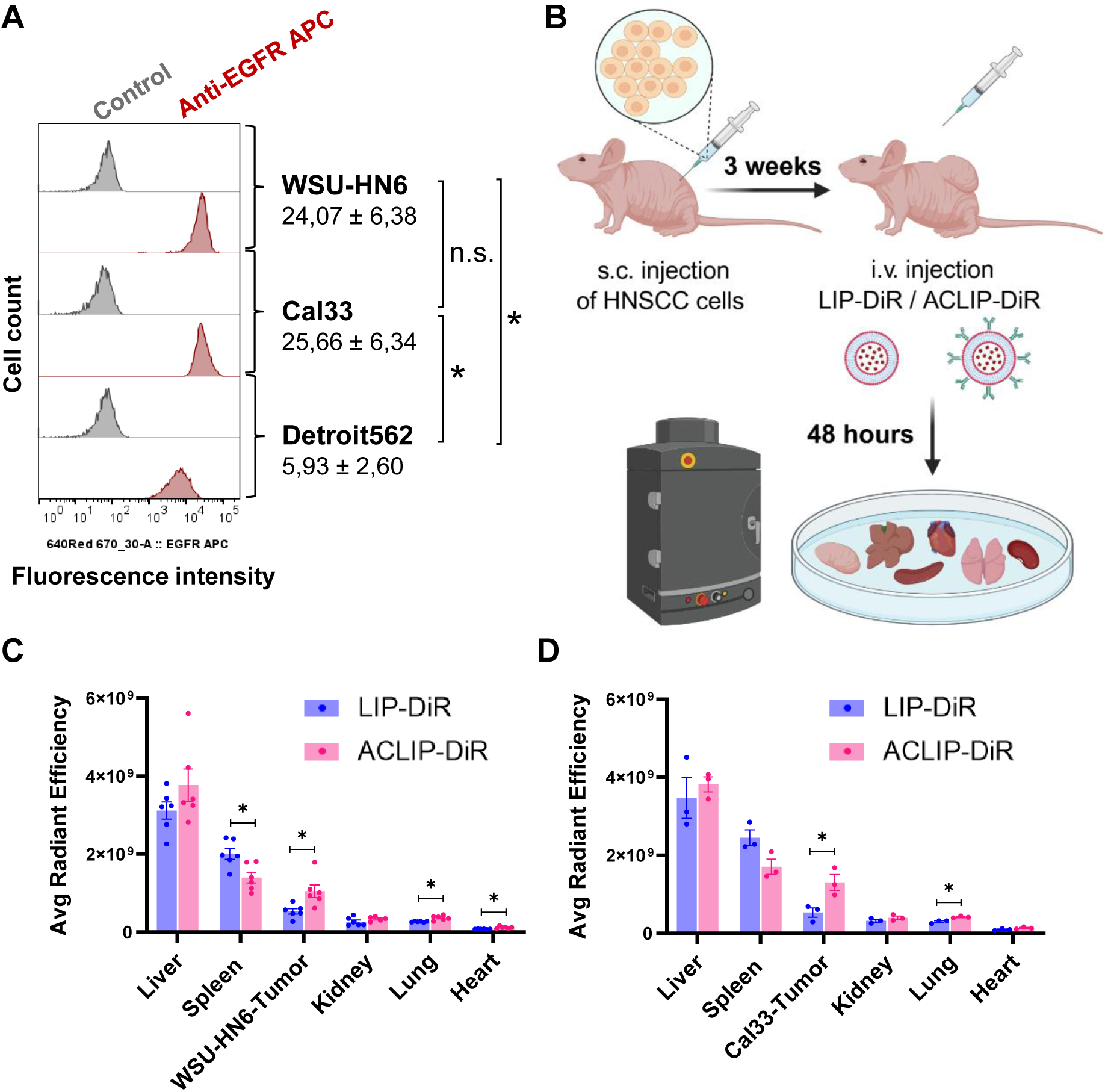
A) Overlay plot of flow cytometry histograms of EGFR expression on the surface of the indicated cell lines (Y axis: cell count; X axis: fluorescence intensity). Data show median fluorescence X1000 ± median absolute deviation for anti-EGFR APC samples (n = 3). One-way ANOVA with Tukey’s corretion for multiple comparisons test. * 0.01 < p ≤ 0.05; n.s., not significant. **B)** Schematics of the murine xenograft model for the analysis of liposome tumor homing. Two million WSU-HN6 or five million Cal33 HNSCC cells were subcutaneously (s.c.) injected into the right flank of immunocompromised nude mice. After 3 weeks, mice were injected intravenously (i.v.) with 100 μL of LIP-DiR or ACLIP-DiR, and *ex vivo* fluorescence was analyzed 48 hours later using an IVIS Lumina In Vivo imaging system. The average radiant efficiency [p/s/cm²/sr] / [µW/cm²] of the indicated organs and tumors from mice injected with **(C)** WSU-HN6 or **(D)** Cal33 cells was measured. Data represent mean ± SEM (n = 6 for WSU-HN; n = 3 for Cal33). Unpaired t-test. Welch’s correction was used when variances differed (F test). Only significant comparisons are indicated: * 0.01 < p ≤ 0.05; the rest are n.s.

After the tumors had developed, mice were injected with the DiR-labeled liposomes. Forty-eight hours later, we analyzed the fluorescence intensity in various organs *ex vivo*. Consistent with the pharmacokinetic behavior of liposomal carriers [53,54], fluorescence accumulated mainly in the liver and spleen, followed by the tumors. Mice that received ACLIP-DiR exhibited higher tumor fluorescence signal than those injected with LIP-DiR in both models (Figure 5C&D), indicating that active targeting enhances the accumulation of liposomes at the tumor site. We observed decreased spleen accumulation of ACLIP-DiR compared to LIP-DiR (statistically significant in the WSU-HN6 model).

### 3.6. Evaluation of free CDDP and ACLIP-CDDP in an EGFR-positive HNSCC xenograft model

Given that CDDP remains a standard therapy in HNSCC and the substantial toxicity associated with its use [55,56], we selected ACLIP-CDDP to evaluate its effectiveness in inhibiting tumor growth in an EGFR-positive HNSCC xenograft model (Figure 6A). Treatment with ACLIP-CDDP, as well as free CDDP, significantly decreased the growth of Cal33 tumors compared to Control (Figure 6B). In our study, tumor size and cell proliferation reduction in mice treated with ACLIP-CDDP was similar to that in the free CDDP treated group (Figure 6B and C).

**Figure 6.**
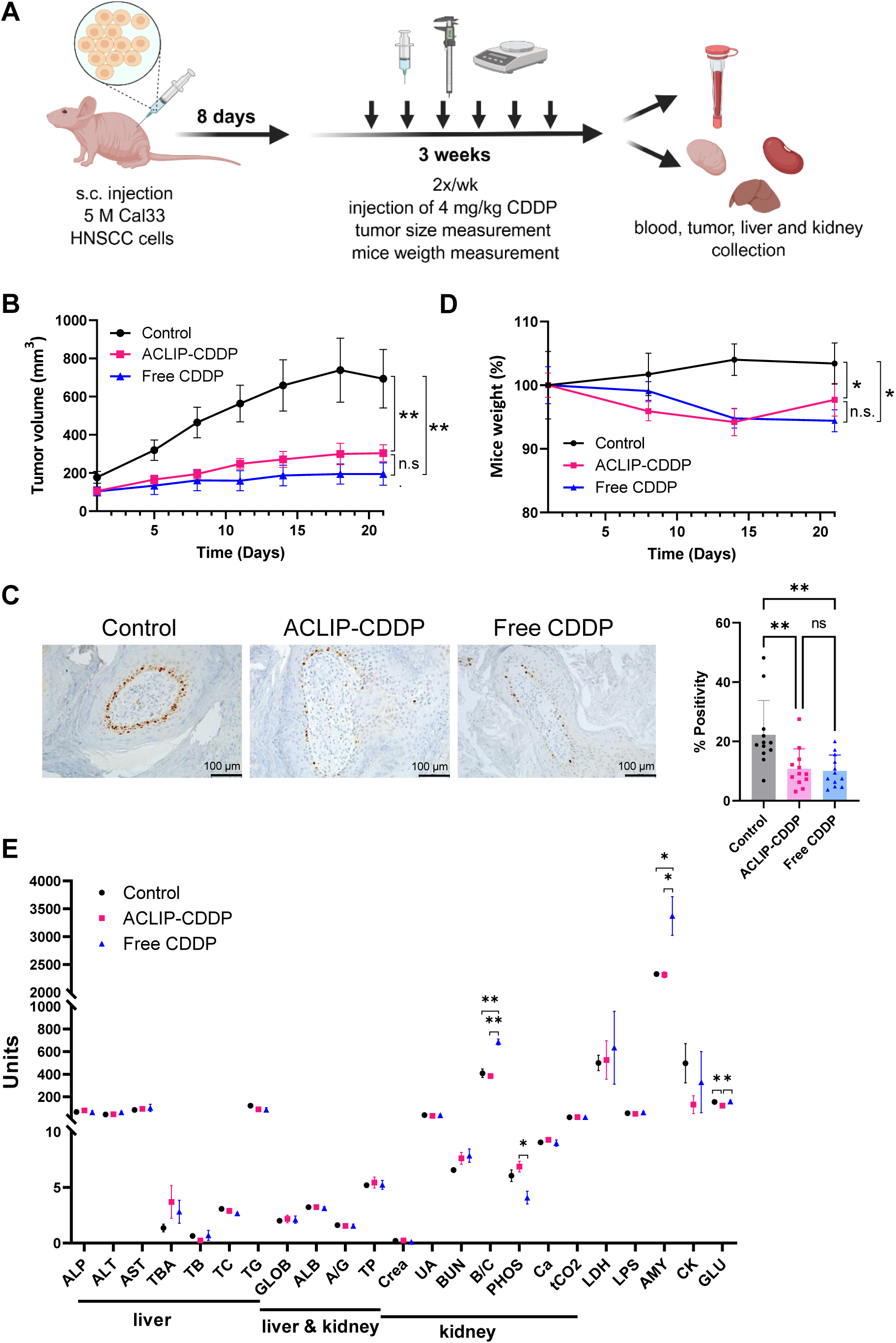
A) Schematic representation for the experimental design of the murine xenograft model for the analysis of tumor growth. Five million Cal33 HNSCC cells were subcutaneously (s.c.) injected into both flanks of immunocompromised nude mice (n = 4 mice/8 tumors per treatment group). Eight days after injection, mice were treated with 4 mg/kg CDDP (free CDDP or ACLIP-CDDP) twice a week for 3 weeks (day 1 to day 21). Tumor volume was measured twice a week and animal weight once a week, for 3 weeks. Blood, tumors and organs were collected at the end of the treatment. **B)** Growth curves of Cal33 xenograft tumors treated with empty ACLIP (Control), free CDDP, and ACLIP-CDDP. Data represent mean ± SEM (n = 8). One-way ANOVA with Tukey’s correction for multiple comparisons test. ** p ≤ 0.01; n.s., not significant. **C)** Representative images of immunohistochemical staining and quantification (right panel) for Ki67 in xenograft tumor tissues from mice treated with the indicated conditions. For the quantification, Ki67-stained slides from four representative images from three different tumors were photographed at 200X magnification. Positively and negatively Ki67-stained cells were counted. Data represent mean ± SEM (n = 12 slices/per treatment group; at least n = 8,500 cells/treatment group). One-way ANOVA with Tukey’s correction for multiple comparisons test. ** p ≤ 0.01; n.s., not significant. **D)** Body weight of mice during the treatment. Data represent mean ± SEM (n = 4). One-way ANOVA with Tukey’s correction for multiple comparisons test. * 0.01 < p ≤ 0.05; n.s., not significant. **E)** Blood samples were collected at the end of the experiment (day 21). The parameters analyzed and the measurement units are as follows. Liver function: ALP (alkaline phosphatase, U/L), ALT (alanine aminotransferase, U/L), AST (aspartate aminotransferase, U/L), TBA (total bile acids, µmol/L), TB (total bilirrubin, mg/dL), TC (total cholesterol, mmol/L), TG (triglycerides, mg/dL). Liver and/or renal function: GLOB (globuline, g/dL), ALB (albumin, g/dL), A/G (albumin/globulin ratio), TP (total protein, g/dL). Renal function: Crea (creatinine, mg/dL), UA (uric acid, µmol/L), BUN (blood urea nitrogen, mmol/L), B/C (BUN/Crea ratio), PHOS (phosphorus, mg/dL), Ca (calcium, mg/dL), tCO2 (bicarbonate, mmol/L). Tissue damage: LDH (lactate dehydrogenase, U/L). Pancreatic function: LPS (lipase, U/L), AMY (amylase, U/L). Muscle damage: CK (creatine kinase, U/L). Diabetes parameters: GLU (glucose, mg/dL). Data represent mean ± SEM (n = 3). One-way ANOVA with Tukey’s correction for multiple comparisons test. * 0.01 < p ≤ 0.05; ** p ≤ 0.01; n.s., not significant.

Mice treated with free CDDP and ACLIP-CDDP experienced a slight (less than 10%) but significant weight loss compared to the Control group. At the end of the experiment on day 21, weight loss was more pronounced in the mice treated with free CDDP than in those treated with ACLIP-CDDP, though this difference did not reach statistical significance (Figure 6D).

The kidney is the primary organ affected by CDDP therapy. Renal toxicity is the main dose-limiting side effect and patients with compromised renal function are often not suitable for CDDP therapy. The analysis of the blood samples collected at the end of the experiment (day 21) (Figure 6E) showed that mice treated with free CDDP had a significantly increased blood urea nitrogen/creatinine (B/C) ratio and decreased level of phosphorus (PHOS) compared to those treated with ACLIP-CDDP. Both of these parameters are associated with impaired kidney function. Similarly, free CDDP-treated mice displayed higher levels of amylase (AMY) compared to the ACLIP-CDDP group. While alterations in pancreatic function are considered the main cause of AMY raise in blood, it has been reported that the levels of this enzyme tend to increase when kidney function declines [57–59].

We did not observe any changes in blood parameters (Figure 6E) and tissue histology (Figure S6 in the Supplementary Material) associated with liver damage. This is of particular importance in mice treated with empty liposomes (Control) and with ACLIP-CDDP, since liposomes tend to accumulate in the liver.

These findings suggest that, under the experimental conditions tested, ACLIP-CDDP did not show superior efficacy in inhibiting tumor growth compared to free CDDP. Nonetheless, encapsulating CDDP in targeted liposomes decreased the signs of renal toxicity, highlighting the clinical relevance of strategies for CDDP encapsulation and tumor targeting.

## 4. Discussion

HNSCC accounts for approximately 90% of all HNCs and is the sixth most common cancer worldwide [1,2]. Systemic toxicity, poor selectivity, and the development of resistance hinder conventional chemotherapy for HNSCC [7,8,13–15]. Targeted therapies for HNSCC remain limited. The anti-EGFR antibody Cetuximab and the anti-PDL-1 antibodies Pembrolizumab and Nivolumab were approved in 2006 and 2016, respectively, for the treatment of advanced or recurrent tumors. However, response rates remain modest, benefiting only about 20% of patients. Small molecule inhibitors are currently under clinical investigation for HNSCC, but none have yet reached clinical approval.

In this context, we explored nanomedicine as a promising strategy to address the limitations of both conventional and emerging targeted therapies in HNSCC, such as suboptimal treatment efficacy and harmful off-target effects [16,17]. These hindrances are especially relevant for treatments based on CDDP, which remain a cornerstone treatment for locally advanced HNSCC [60,61]. Therapies with novel small-molecule inhibitors, such as the PI3K/AKT/mTOR pathway inhibitor Alpelisib, in clinical studies for this tumor type [62–64], could also benefit from these approaches.

Converting conventional drugs into nanomedicine formulations offers other therapeutic advantages, including controlled and sustained drug release, improved pharmacokinetics and biodistribution, and the potential to overcome drug resistance [65]. Moreover, nanotechnology enables the transformation of conventional therapies into targeted treatments through nanocarrier design. In this study, we actively targeted liposomes towards tumor cells using anti-EGFR antibodies. The frequent overexpression of EGFR in various epithelial malignancies supports this strategy [66]. In HNSCC, 90% of tumors overexpress EGFR mRNA [24], and the protein levels correlate with reduced overall and progression-free survival [25]. Fluorescence particles, including nanoparticles and antibody conjugates, targeting this receptor have been proposed as tools to aid the detection of surgical tumour borders in intraoperative fluorescence-guided surgery [67–69].

We successfully formulated passive and active-targeting liposomes based on optimized protocols previously developed and validated by our research group (Scheme 1) [35–37]. DDS were obtained *via* solvent-injection methods, while active targeting was achieved through carbodiimide-mediated conjugation, a widely adopted approach owing to its cost-effectiveness, simplicity, and high coupling efficiency [45,46]. Liposomal formulations were extensively characterized based on hydrodynamic radius, polydispersities, morphology, encapsulation efficiencies, stability over time, and drug release profiles. The formulations exhibited a characteristic spherical morphology and demonstrated stability, as well as controlled drug release profiles. Evidence of successful Cetuximab conjugation was supported by zeta potential shifts and modifications of the liposomal surface observed by TEM imaging.

*In vitro* preclinical assessment showed that both passive and active-targeting liposomes are promising DDS for Alpelisib, as they demonstrated equipotent antiproliferative activity in three HNSCC cell lines with distinct PIK3CA profiles (activating mutation, gene amplification, and wild-type). To our knowledge, this is the first report of Alpelisib encapsulation in lipid nanoparticles. These findings encourage further *in vivo* preclinical studies, where factors such as tumor microenvironment, vascularization, immune response, and tissue-specific drug accumulation play crucial roles. While achieving therapeutic doses of Alpelisib encapsulated into active-targeting liposomes suitable for *in vivo* studies remains a work in progress, we successfully evaluated formulations based on CDDP, the mainstay chemotherapy in HNSCC, to assess active targeting of drug-loaded liposomes both *in vitro* and *in vivo*. Similar to Alpelisib, passive and active-targeting CDDP-liposomes displayed comparable *in vitro* antiproliferative responses in three HNSCC cell lines with varying sensitivity to CDDP.

To evaluate the *in vivo* effect of the active-targeting CDDP nanomedicine, we performed human xenografts using the HNSCC-derived Cal33 cell line, which expresses EGFR on the cell surface. Unexpectedly, the active-targeting CDDP liposomes did not outperform free CDDP in inhibiting tumor growth. This outcome may be attributable to the physicochemical properties of CDDP, a prodrug that becomes highly hydrophilic upon activation through aquation [70–72]. As suggested by the *in vitro* release profile, the increased hydrophilicity of activated CDDP could facilitate its premature release from the liposomal carrier. *In vivo*, this activation may enhance CDDP diffusion into the bloodstream, thereby reducing the amount of intact formulation reaching the tumor site.

We conducted *in vivo* imaging studies to elucidate whether the outcome in our *in vivo* preclinical evaluation was attributable to limitations of the active-targeting approach or constraints associated with the liposomal delivery platform. In two EGFR-positive xenograft models, active-targeting liposomes loaded with the near-infrared fluorescent DiR dye showed higher tumor accumulation and better tumor-to-spleen ratio compared to the passive liposomes. These findings suggest that the lack of therapeutic superiority in tumor growth inhibition of the active-targeting nanomedicine compared to the free drug may stem from drug-specific limitations, which may lead to premature release before tumor accumulation, rather than failure of the targeting system.

Encapsulating CDDP is one of the most promising and actively investigated drug delivery strategies for HNSCC therapy. Others have reported an inhibitory effect of CDDP on tumor growth using both passive and active-targeting strategies. However, these reports are based on co-encapsulating CDDP with other drugs or on intratumoral drug delivery strategies [30–34]. Similar to our findings, using an immunocompetent mouse model, Lan et al. [31] did not find significant differences in tumor growth inhibition and immune profile between two systemic treatments based on the combination of anti-PD-1 with free CDDP and anti-PD-1 with liposome-encapsulated CPPD.

Previous clinical trials using liposomal CDDP formulations have shown promising safety profiles and reduced toxicity [73–76]. In HNSCC, a phase I study using Stealth® liposomal CDDP with radiotherapy reported minimal systemic toxicity, and no instances of nephrotoxicity, ototoxicity, neurotoxicity, or dose-limiting toxicities observed at the highest doses tested [74]. Lipoplatin (pegylated liposomes encapsulating CDDP) decreased the typically CDDP-associated adverse effects in a phase III pilot trial in patients with non-small cell lung cancer or neuroblastoma [73]. Both studies revealed that CDDP-loaded DDS reduce renal and hematological toxicity to a clinically relevant extent.

Consistent with these findings, our blood analysis also suggested favorable nephrotoxicity profiles for the active-targeting nanomedicine compared to the free drug. Importantly, we did not observe hematological or histological signs of liver damage despite the typical accumulation of liposomes in this organ.

Importantly, systemic exposure to our active-targeting liposome-released drug is expected to remain lower than that of free CDDP, as indicated by both the tumor accumulation observed with the liposomal formulation and reduced renal toxicity in the blood profile of the active-targeting CDDP-liposome treated group. In this context, exploring higher dosing regimens could further enhance antitumor efficacy, since the improved therapeutic index of the liposomal system allows for potential dose escalation without increasing systemic adverse effects. This hypothesis is further supported by the clinical data reported for other liposomal CDDP formulations, as discussed above.

## 5. Conclusions

In this study, we successfully developed passive and active-targeting liposomal formulations of CDDP and Alpelisib for the treatment of HNSCC. Despite their therapeutic relevance, significant dose-related toxicities limit the use of both drugs. Our *in vitro* studies demonstrated that passive and active formulations maintained antiproliferative activity comparable to that of the free compounds, supporting the feasibility of these delivery strategies for HNSCC treatment. Notably, *in vivo* studies showed that actively targeted CDDP liposomes achieved similar levels of tumor suppression as the free drug with a more favorable nephrotoxicity profile. This outcome is likely attributable to the physicochemical properties of CDDP, since fluorescent-tracking experiments confirmed enhanced tumor accumulation of actively targeted liposomes, demonstrating the potential of active-targeting nanocarrier systems for improving drug delivery in HNSCC.

Despite the limitations observed, this work lays a foundation for future studies aimed at optimizing active-targeting nanoplatforms that can help overcome the toxicity-related drawbacks of current therapies for HNSCC.

## Funding

Iván Bravo and Carlos Alonso-Moreno’s lab is supported by the Ministerio de Ciencia, Innovación y Universidades and the Agencia Estatal de Investigación, Spain, MICIU/AEI/10.13039/501100011033 (grants CPP2021-008597 and RED2022-134287-T), grant 2021-GRIN-31240 funded by Universidad de Castilla-La Mancha, JCCM and EU through INNOCAM and Fondo Europeo de Desarrollo Regional, FEDER (project number SBPLY/21/180501/000050), and ACEPAIN Foundation. F. de Andrés’ lab is supported by Spanish Ministry of Science and Innovation (grant number PID2022-138761NB-100) and the JJCC Castilla-La Mancha (grant number JCCM SBPLY/21/180501/000188). The work developed by CIEMAT/i+12/CIBERONC was funded by projects PI21/00208 and CB16/12/00228 from the Instituto de Salud Carlos III and co-funded by the European Union (EU). C.M-H. was supported by a predoctoral research grant from Comunidad Autónoma de Madrid (ref PIPF-2023_SAL-GL-29800). C.H-I. was supported by a European Social Fund Plus (ESF+) co-funded grant from Comunidad Autónoma de Madrid (ref PEJ-2023-AI_SAL-GL-27408).

## Declaration of generative AI and AI-assisted technologies in the manuscript preparation process

During the preparation of this work the author(s) used the free version of Grammarly in order to obtain **grammar and spelling suggestions**. We used the “Review suggestions” tool, and the author(s) reviewed and edited the content as needed and take(s) full responsibility for the content of the published article.

## Notes

### Competing Interest Statement

The authors have declared no competing interest.

