## Supplementary Figure for "Active Liposomal Targeting for Head and Neck Squamous Cell Carcinoma Treatment"

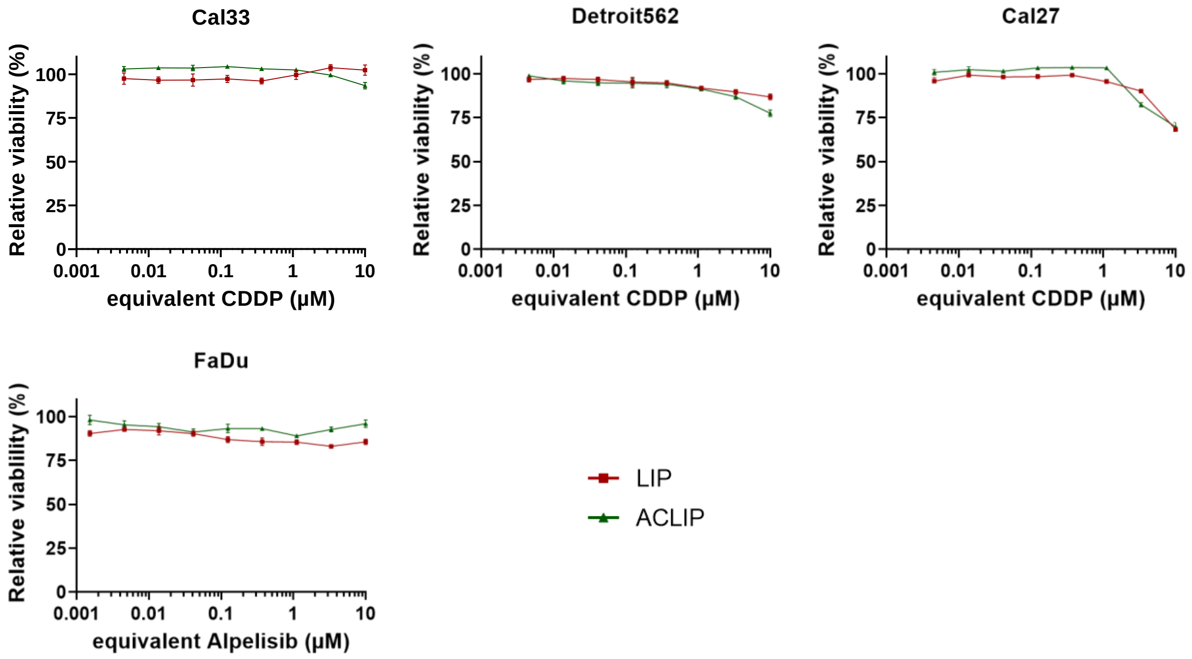


**Figure S3.** Cell viability of the indicated cell lines treated with increasing concentrations of empty LIP and ACLIP. The data are mean ± SEM (n = 3). The amounts of LIP or ACLIP at each point are equivalent to those of the corresponding CDDP or Alpelisib concentration. CDDP: Cisplatin; LIP: liposome; ACLIP: Cetuximab-conjugated liposome.

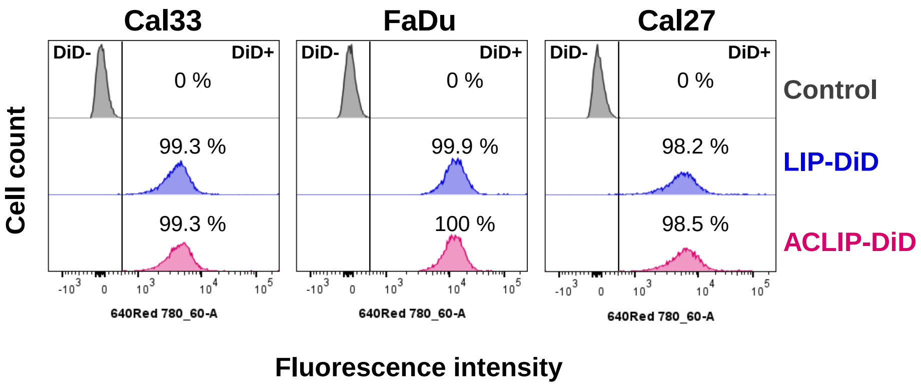


**Figure S4.** Overlay plots of flow cytometry histograms of the indicated cell lines incubated for 30 min in the presence of LIP-DiD (blue peaks) or ACLIP-DiD (magenta peaks). Y axis: cell count; X axis: DiD fluorescence intensity. Background fluorescence in the absence of DiD-loaded liposomes is indicated by the grey peaks. Numbers indicate % of DiD positive cells. LIP-DiD: liposome encapsulated DiD; ACLIP-DiD: Cetuximab-conjugated liposome encapsulated DiD.

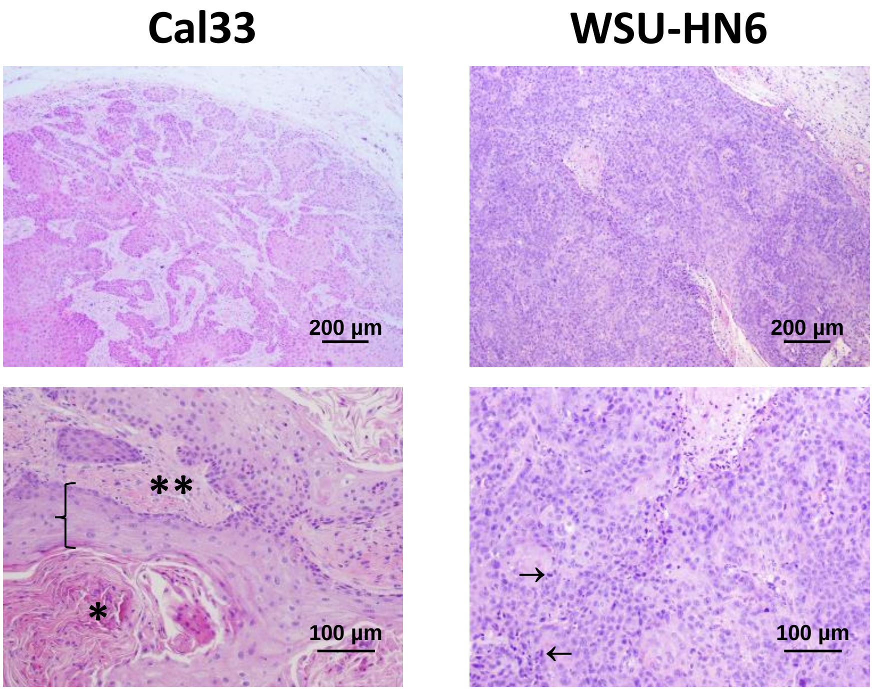


**Figure S5.** H&E staining of 5 µm sections of paraffin-embedded xenograft tumors generated with the indicated HNSCC cell lines. Well-differentiated Cal33-tumors are more eosinophilic and characterized by the presence of keratin pearls (*), a fibrous stroma (**), and features of squamous differentiation (curly bracket). Poorly-differentiated WSU-HN6 tumors are typically highly disorganized and have lost the cytological features characteristic of squamous differentiation. They lack keratin pearls and display a high mitotic rate with frequent atypical mitoses (→).

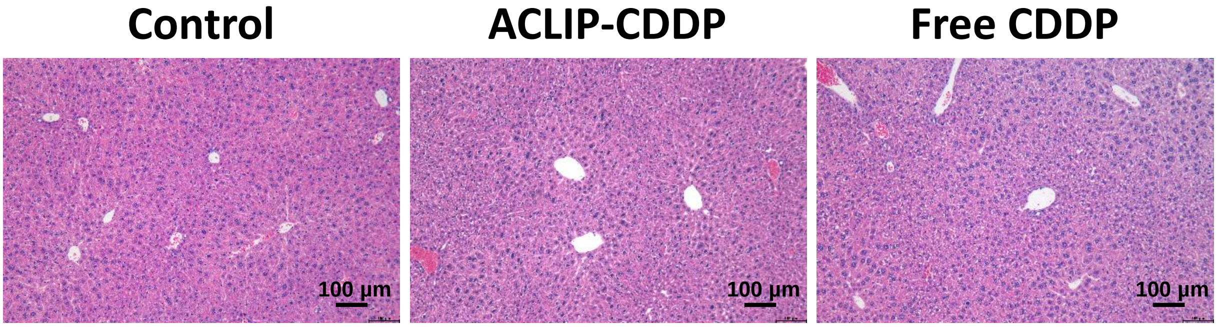


**Figure S6.** H&E staining of 4 µm sections of paraffin-embedded livers from the xenograft HNSCC model collected at the end of the experiment (day 21). Hepatic structure is conserved, with hepatocytes radiating from the central veins. No significant inflammation, necrosis, or fibrosis is observed.
